# Multi-omics decipher the molecular mechanisms driving high-lipid production in an artificially-evolved *Chlamydomonas* mutant

**DOI:** 10.1101/2025.08.05.668592

**Authors:** David R. Nelson, Amphun Chaiboonchoe, Weiqi Fu, Basel Khraiwesh, Bushra Dohai, Ashish Jaiswal, Dina Al-Khairy, Alexandra Mystikou, Latifa Al Nahyan, Amnah Salem Alzahmi, Layanne Nayfeh, Sarah Daakour, Matthew J O’Connor, Mehar Sultana, Khaled Hazzouri, Jean-Claude Twizere, Kourosh Salehi-Ashtiani

## Abstract

Enhancing lipid accumulation in microalgae is critical for commercial viability but often compromises growth. We previously identified an artificially evolved Chlamydomonas reinhardtii mutant (H5) that retains wild-type growth (CC-503) while producing significantly more lipids. Here, we present multi-omic analyses that elucidate the molecular basis of this phenotype. Whole-genome sequencing revealed over 3,000 mutations in H5, including 45 in protein-coding genes (e.g., phosphofructokinase, acyl-carrier protein, glycerol kinase). Six corresponding CLiP insertion mutants also showed elevated lipid content. Transcriptomics revealed upregulation of key genes for glycolysis, nutrient uptake, and proliferation (e.g., pyruvate carboxylase, carbonic anhydrase) under nutrient-replete conditions. Metabolomics identified a striking increase in malonate, a metabolite that supports fatty acid synthesis and cell proliferation. Epigenomic profiling showed hypomethylation in triacylglycerol (TAG) biosynthesis genes and hypermethylation in energy balance regulators. Together, these data suggest that accelerated glycolysis and streamlined metabolism drive lipid accumulation in H5 without compromising growth. Our findings provide a blueprint for engineering high-lipid microalgal strains for industrial applications.

**HIGHLIGHTS:**

- High-lipid *Chlamydomonas* mutant (H5) exhibits cancer-like metabolism: pseudo-hypoxia and nutrient deprivation response
- Multi-omics reveals 45 high-impact mutations synergistically enhance lipid production in H5
- Six CLiP mutants of H5-disrupted genes showed significantly increased lipid content
- Malonate levels increased 10-fold in H5, indicating altered mitochondrial function
- H5 upregulates glycolytic genes while maintaining wild-type growth rates
- Transcriptomes from H5 and CC-503 converge after nitrogen deprivation despite replete-state differences
- H5 shows altered lipid composition with increased TAG diversity, decreased DAGs
- Epigenomic profiling reveals 14,720 differentially methylated transcribed regions in H5

## INTRODUCTION

Algae are a promising platform for sustainable production of biofuels, pharmaceuticals, and other high-value products, due to their rapid growth and capacity to thrive on non-arable land [1–6]. Unlike land crops, they support continuous, season-independent cultivation [7, 8] and can be grown in regions unsuitable for agriculture, offering an alternative to deforestation-linked crops like oil palm [9–12].

However, lipid accumulation competes with cell proliferation for energy and carbon, creating a trade-off that limits productivity [13]. Fatty acids can be routed either to membrane synthesis or to intracellular lipid bodies—the latter being most relevant for industrial applications [13]. Wild-type strains typically exhibit low lipid content, and while transgenic efforts have achieved incremental improvements, they often face instability or reduced growth. Overcoming the growth–lipid tradeoff is essential for profitable algal oil production, yet current engineering approaches appear insufficient for sustained gains [14].

*Chlamydomonas reinhardtii* has arguably been the most popular and well-studied green microalga to date. *Chlamydomonas*, although not oleaginous by definition, is widely used to study microalgal lipid metabolism because it, like many other microalgae, accumulates triacylglycerols under various conditions [15]. Our study focuses on a high-lipid *C. reinhardtii* mutant (named ‘H5’) with multiple mutations. We aim to understand the underlying mechanisms in artificial laboratory evolution (ALE) to produce a synthetic microbial chassis suitable for profitable biotechnological lipid production in microalgae. Specifically, we target microalgae growing exponentially, working towards a continuous production system for microalgal lipids.

Our previous work [16, 17] indicated that UV mutagenesis, combined with iterative rounds of selection, could be an effective strategy to increase lipid accumulation in microalgae. The CC-5163 mt+ mutant (H5, found at https://www.Chlamycollection.org) was generated through a series of UV mutagenesis and selection cycles from the starting strain CC-503 [16]. Optimizing oil production using this strategy requires sorting and analyzing many algal isolates based on their triacylglycerol (TAG) contents [16, 17]. We found that H5 has enhanced lipid accumulation compared to its progenitor strain, CC-503, as determined by flow cytometry and fluorescence microscopy (Fig. 1). Flow cytometry data for cell populations from different growth phases showed more lipid accumulation in nutrient-replete conditions for H5 (Fig. 1). The mutant H5 made 3.2-fold more lipid than its parental strain CC-503 in exponential phase growth (OD_600_ = 0.4 [16]). Consistent with flow cytometry measurements, confocal microscopy of cells stained with lipophilic dyes showed that H5 has more lipid droplets (LDs) with brighter BODIPY fluorescence than CC-503 (Fig. 1). Under mixotrophic growth conditions, H5 had a similar growth rate compared to its parental counterpart (Fig. S1), which indicates steady-state regulatory shifts in lipid metabolism in the mutant.

**Figure 1.**
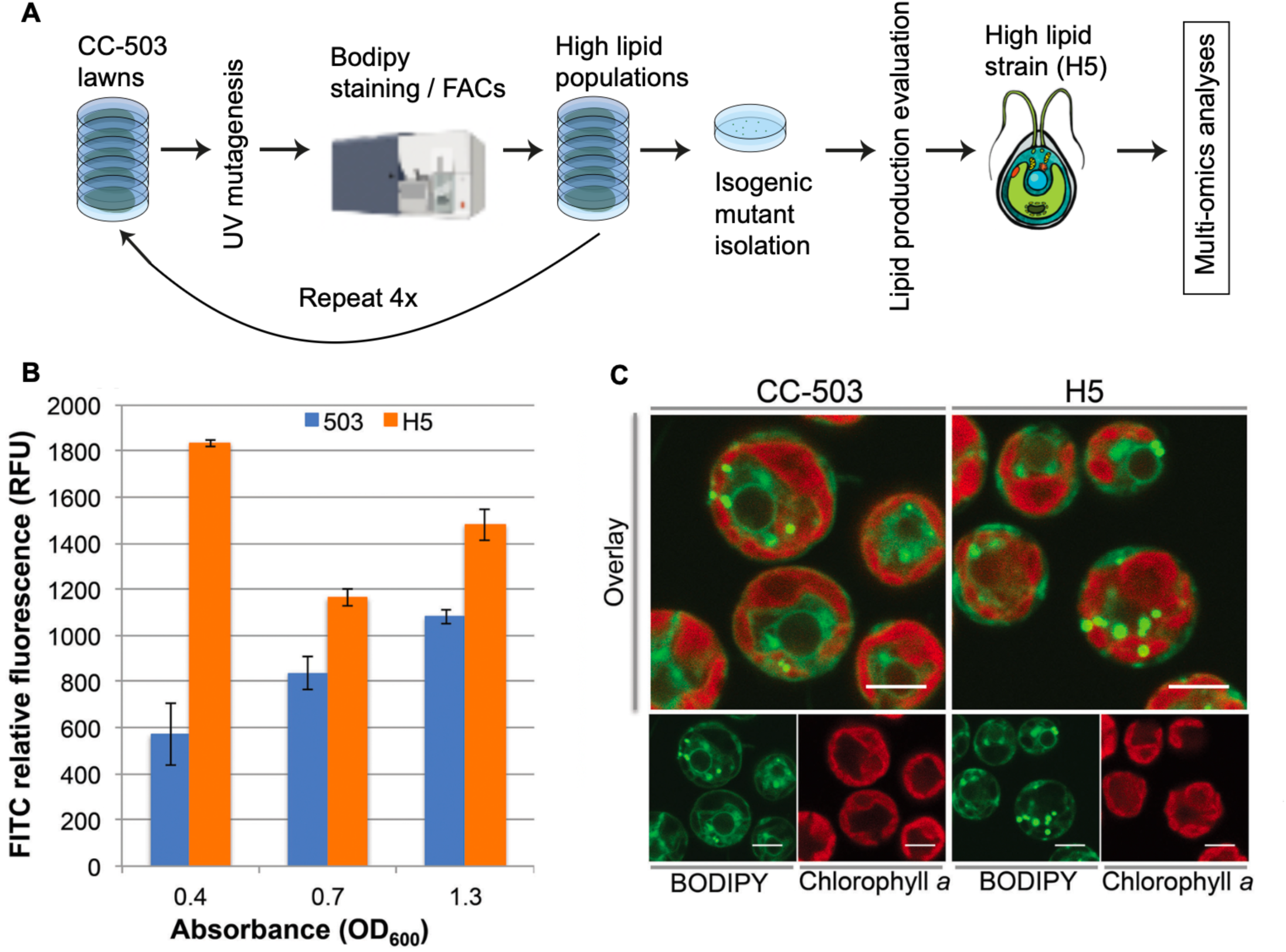
Phenotype and genotype of H5 compared to wild-type (CC-503). **(A)** Schematic diagram showing the overall experimental procedure to generate high-lipid *C. reinhardtii* strains (adapted from Abdrabu et al. [16, 17] and Sharma et al.[17]). **(B)** Flow cytometry-based fluorescence intensity measurements for BODIPY 505/515-stained *C. reinhardtii* cells in different growth phases indicated by their optical density. Standard error is shown. H5 cells accumulated significantly more lipids in the exponential growth phase (OD_600_ = 0.4). **(C)** Confocal fluorescence microscopy of BODIPY 505/515-stained CC-503 and H5 strains contrasted by chlorophyll autofluorescence during the exponential growth phase. Scale bar indicates 5 µm.

## RESULTS AND DISCUSSION

A wide variety of mutagenic events, including single nucleotide polymorphisms, deletions, and inversions, can occur in eukaryotic cells in response to UV irradiation, followed by photo repair inactivation [18] during cell division [19–22]. We interrogated the underlying genetic and metabolic events that have resulted in lipid accumulation without compromising growth in H5. Whole-genome resequencing of the mutant strain revealed multiple UV-induced mutations [19–22], including single nucleotide polymorphisms (SNPs), insertions, deletion, and multiple nucleotide polymorphisms (MNPs) after aligning the obtained genomic reads to the reference genome of the parent strain CC-503. Transcriptomic and metabolomic analyses revealed altered metabolism in H5, including glycerolipid and phospholipid pathways, increased expression of glycolytic enzymes and products, and a pseudo-nutrient deprivation response characterized by the upregulation of carbon, nitrogen, molybdenum, iron, and oxygen deprivation responsive genes.

### Whole-genome sequencing identifies variants in lipid biosynthesis pathways

We sequenced the genome of two isogenic H5 populations to characterize UV-induced mutations and their frequency and position relative to protein-coding regions. Cells grown from two independent colonies were sequenced to increase confidence in mutational event identification. After the mapping of filtered Illumina HiSeq 2500 reads to the reference CC-503 genome [23], SnpEff [24] was used to find the impacts of the variant on known genes based on the detected variants, including single nucleotide polymorphism (SNPs), insertions, deletions, and multiple nucleotide polymorphisms (MNPs). Impacts on encoded protein functions were graded as high, moderate, or low. We found 5,341 variants, including 3,984 SNPs, 647 insertions, and 710 deletions (Fig. 2 and Table S3). Most identified variations were associated with regions upstream and downstream to genes (approximately 30%; Fig. 2, Table S2). The most frequent base change was C –> T (n = 690), followed by G –> A (n = 632), A –> G (n = 493), and T –> C (n = 470) base changes (Fig. 2). We note that C –>T transitions are common signatures of pyrimidine dimer repair after UV exposure in many organisms [25]. Our data suggest that *C. reinhardtii* also follows this trend. However, the ratio of C –> T transitions introduced compared to other mutation classes likely depends on the time spent in the dark to prevent photoreactivation [26–28] and the UV exposure duration (multiple nucleotide track polymorphisms) and intensity (double-strand breaks, insertions and, deletions) in the experimental conditions.

**Figure 2.**
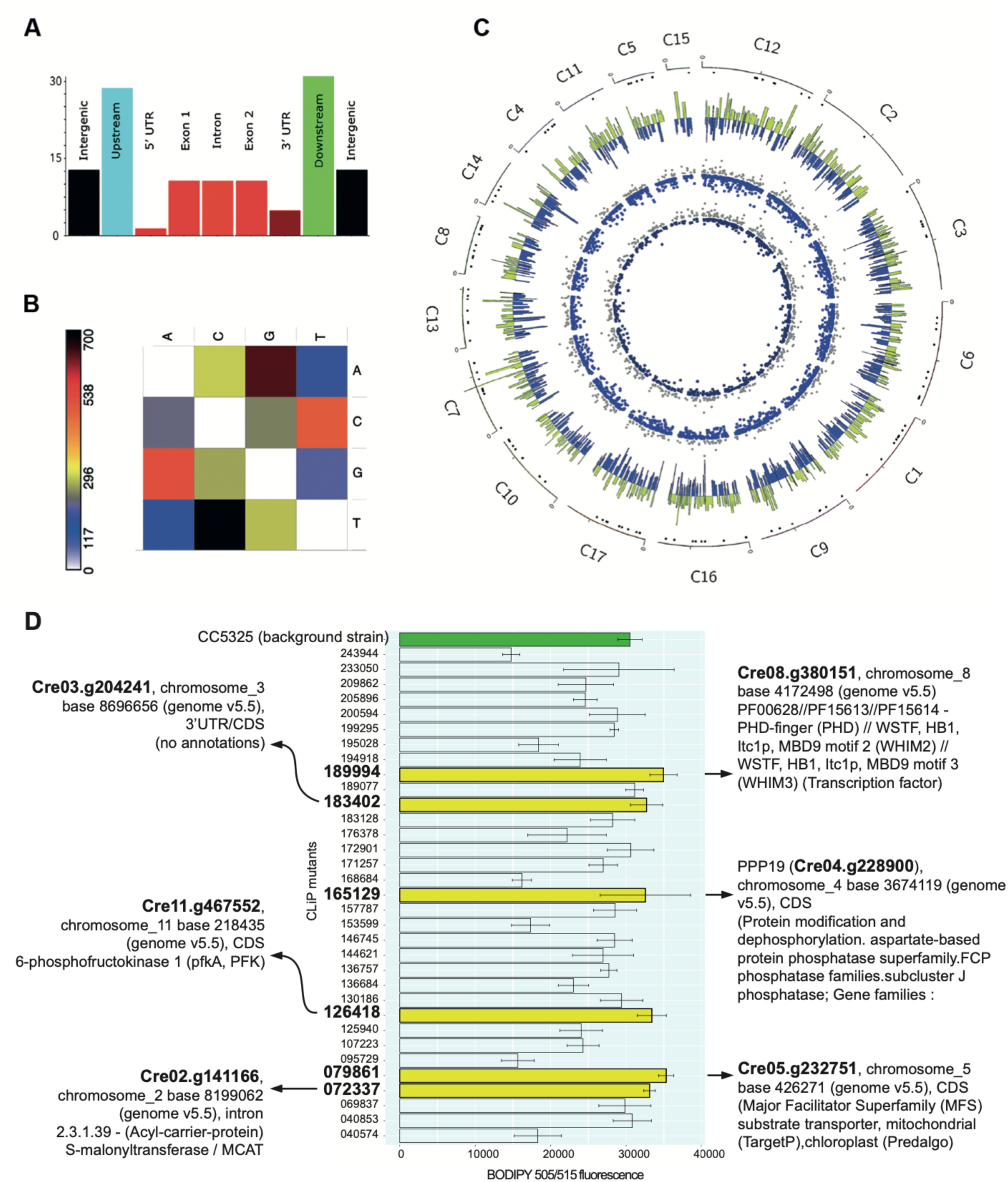
Single nucleotide polymorphisms (SNPs) in the mutant H5 compared to the parental CC-503. **(A)** Nucleotide variants in H5 compared to CC-503 as discovered by SnpEff [24] from genomic sequencing data (n = 2 for each strain, see Table S1). These variants were predicted to impact the function of their respective genes significantly. Several genes involved in metabolic pathways had frameshift mutations. **(B)** A summary of total base changes for UV-induced mutations in H5. The majority of mutations observed were C –> T transitions, typical products of UV-induced pyrimidine dimer repair.[33] **(C)** Circos (circos.ca) plot showing genomic (SNPeff-predicted mutations) and transcriptomic (RNAseq) data from H5 compared to CC-503 (see also Tables S1 and S3). The outer track indicates genomic mutations, and the inner tracks indicate significant (*q < 0.05)* up and down-regulated genes in the strains for the days sampled. Numbered from outside to inside: 1, dots under chromosome axes represent genes with ‘high-impact’ mutations; 2, yellow histogram facing outwards shows genes upregulated in H5 on Day 0; 3, blue histogram facing inwards shows genes down-regulated in H5 on Day 0; 4, grey scatter plot facing outwards shows genes upregulated in H5 after 24 hours of nitrogen deprivation (performed as in Wang et al., [34]; Day 1); 5, blue scatter plot facing inwards shows genes down-regulated in H5 on Day 1; 6, magenta scatter plot facing outwards shows genes upregulated in H5 after three days of nitrogen deprivation (Day 3); 7, cobalt scatterplot facing inwards shows genes down-regulated in H5 on day 3. In addition to the chromosomes shown in the above plot, several genes were up– and down-regulated in the non-chromosomal genomic scaffolds (see Fig. S2). **(D)** Comparison of lipid accumulation in CLiP [30, 31] strains. BODIPY 505/515 fluorescence of stained cells was determined with an excitation wavelength of 485 nm and an emission wavelength of 515. BODIPY fluorescence was normalized by OD_600_ (x-axis). Standard error is shown. The details of these CLiP mutants are in Table S1 and at Chlamylibrary.org. Six mutants had significantly higher lipid accumulation than CC-503, implying potential roles for their disrupted genes in producing the H5 high-lipid phenotype.

Notably, 45 genes had ‘high-impact’ mutations (Table S1). These mutations, detected by SnpEff [24], were predicted to substantially influence the encoded protein’s function. Here, ‘high-impact’ mutations comprised mutations predicted to substantially perturb the coding sequence or splicing regions of the gene, including missense, frameshift, splice site alteration, and deletion/insertion mutations.

To experimentally determine if the identified genes with ‘high-impact’ mutations could affect lipid accumulation in *C. reinhardtii*, we tested mutant strains from the *Chlamydomonas* Library Project (CLiP) collection [29–32] having insertional mutations in the corresponding genes. Ten genes with putative roles in growth and lipid production were included in the 45 high-impact mutations (Table S1). Thirty-three out of the total of 45 UV-induced mutations in H5 had corresponding knock-in mutants from the CLiP collection [31]. These strains were obtained from www.Chlamycollection.org and tested for increased lipid production using a BODIPY 505/515 fluorescence FACs approach (Table S2). Of these, six strains showed significantly more lipids than the original CC-5325 strain (p < 0.05 in a two-tailed *t* test, Fig. 2, Table S2). In contrast, the other 28 strains showed equal or decreased lipid accumulation. These results suggested that increased lipid accumulation in H5 is, in part, a consequence of the synergistic effects of these mutations.

### Transcriptomes of H5 and CC-503 become similar after nitrogen deprivation

We interrogated the transcriptomes of H5 compared to WT in nutrient-replete and nitrogen-deprived conditions to examine differences in their transcriptomes regarding metabolic regulation and to decipher whether H5 displays a normal nitrogen-deprivation response (Table S3). Transcriptomic sequencing (RNAseq) for CC-503 and H5 cultures in triplicates was performed at 0, 1, and 3 days of nitrogen starvation (see Table S3). Mid-log phase (OD_600_ = 0.4) cells were harvested for the Day 0 timepoint. To define the contributions of deprived and nutrient-replete states towards lipid accumulation in H5 and CC-503, transcriptome analyses were performed at zero, one, and three days post-inoculation in Cuffdiff [35] (Tables S2 and S3). The differentially expressed genes (DEGs) were screened at false discovery rate (FDR)-corrected *p* values (i.e., *q* values) ≤ 0.05 (Table S3). At Day 0, 11,126 and 10,554 genes were expressed in CC-503 and H5 at FPKM > 1.0. Of these expressed genes, 10,094 were shared between the two strains, 1,032 were unique in CC-503, and 460 were unique in H5. Differentially expressed genes from Day 0 (starting point, non-starvation control) and Day 1 (deprived for 24 H) comprised 1,798 and 1,727 upregulated transcripts and 1,296 and 1,602 down-regulated transcripts.

On Day 3 (deprived for 72 H), H5 showed a similar expression profile as CC-503, as only 213 genes were up-regulated and 276 genes were down-regulated (Fig. 3 and Table S3). Thus, H5 exhibits drastically different transcriptome profiles in nitrogen-replete conditions but has transcript expression patterns closer to CC-503 after nitrogen deprivation.

**Figure 3.**
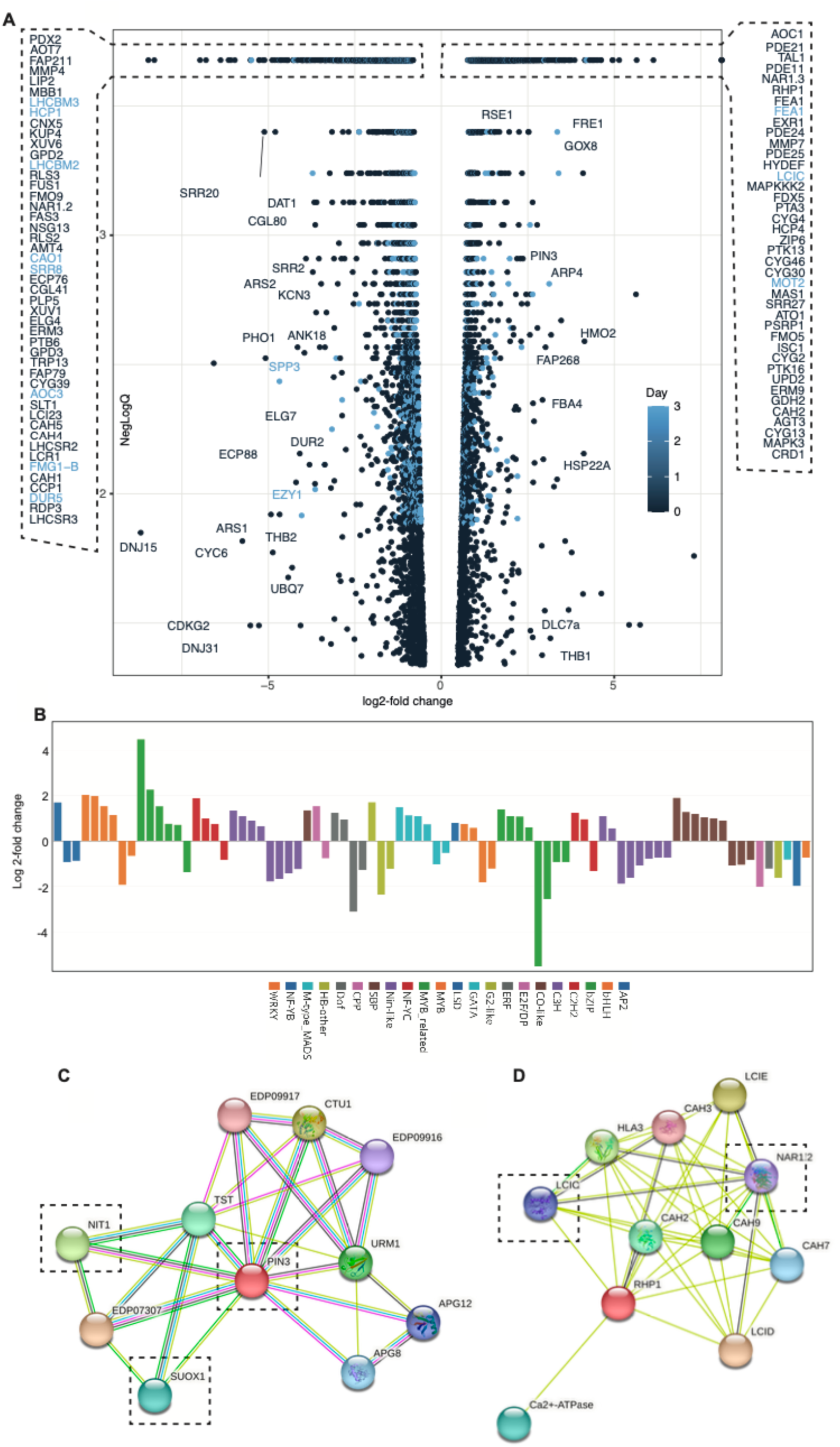
Time-series transcriptomics resolves expression differences in H5 in nitrogen-replete and deprived conditions. (**A**) DEGs between CC-503 and H5 during exponential growth (OD_600_ = 0.4) phase under nutrient-replete (Day 0) and nitrogen-deprived (Day 3) conditions (Table S3). Gene expression in mutants is displayed as a log 2-fold change in H5 compared to CC-503. Log 2-fold changes in each transcript (based on normalized FPKM values) against FDR-corrected *p* values (*q* values). (**B**) Transcription factor landscape of H5 compared with CC-503 at Day 0. Many TFs with roles in cell growth and proliferation were upregulated in H5 (see Table S3). (**C**) STRING (https://string-db.org/) functional association networks of highly-upregulated H5 genes. The central PIN3 is a parvullin homolog of *Arabidopsis* auxin transporters [39]. Red lines represent the presence of fusion evidence; green lines represent neighborhood evidence; blue lines represent co-occurrence evidence; purple lines represent experimental evidence; yellow lines represent text mining evidence; light blue lines represent database evidence, and black lines represent co-expression evidence (see also http://version10.string-db.org). SUOX1 and NIT1 are key enzymes in sulfur and nitrogen metabolism respectively, with SUOX1 oxidizing sulfite to sulfate and NIT1 reducing nitrate to nitrite in *Chlamydomonas*. Their dysregulation could trigger lipid accumulation through nutrient stress signaling and metabolic reallocation of energy resources. (**D**) The carbonic anhydrase (CAH2, Cre04.g223050, log 2-fold change = 2.40148 *q =* 0.000210344, Table S3) was highly upregulated in H5. It is functionally linked (STRING) [40, 41] to the low-carbon responsive proteins LCIC and LCID and a nitrate transporter (NAR). Its expression suggests that H5 has higher carbon and nitrogen assimilation rates than CC-503.

Significant DEGs (Table S3) were classified into functional groups as in Gargouri et al. 2015 [36]. Photosynthesis, TCA cycle, and carbon fixation pathway, as well as multiple key nutrient deficiency-responsive genes (e.g, NIT1, SUOX1, see Fig. 3C-D), were upregulated in H5 compared to CC-503 (Figure 3, Table S3). Several enzymes with predicted high-impact mutations, including a glycerol kinase and a TAG lipase (Table S1), were found to have very high expression levels (Table S3). Forty-six out of 70 lipid metabolism genes were upregulated in H5. These genes were involved in fatty acid, phosphatidylglycerol (PG), and acetyl-CoA synthesis. Also, 40 out of 43 genes involved in photosynthesis were upregulated, including five dominant LHCII complex genes (Table S3). Two mitochondrial carbonic anhydrases (CAH4/5), a cyclin-dependent kinase, pherophorin, and two transporter genes (ABC and anion transporter) were among the top down-regulated genes (Table S3). These genes are likely involved in the dysregulated mitochondrial metabolism predicted in our metabolomic results (Fig. 4).

**Figure 4.**
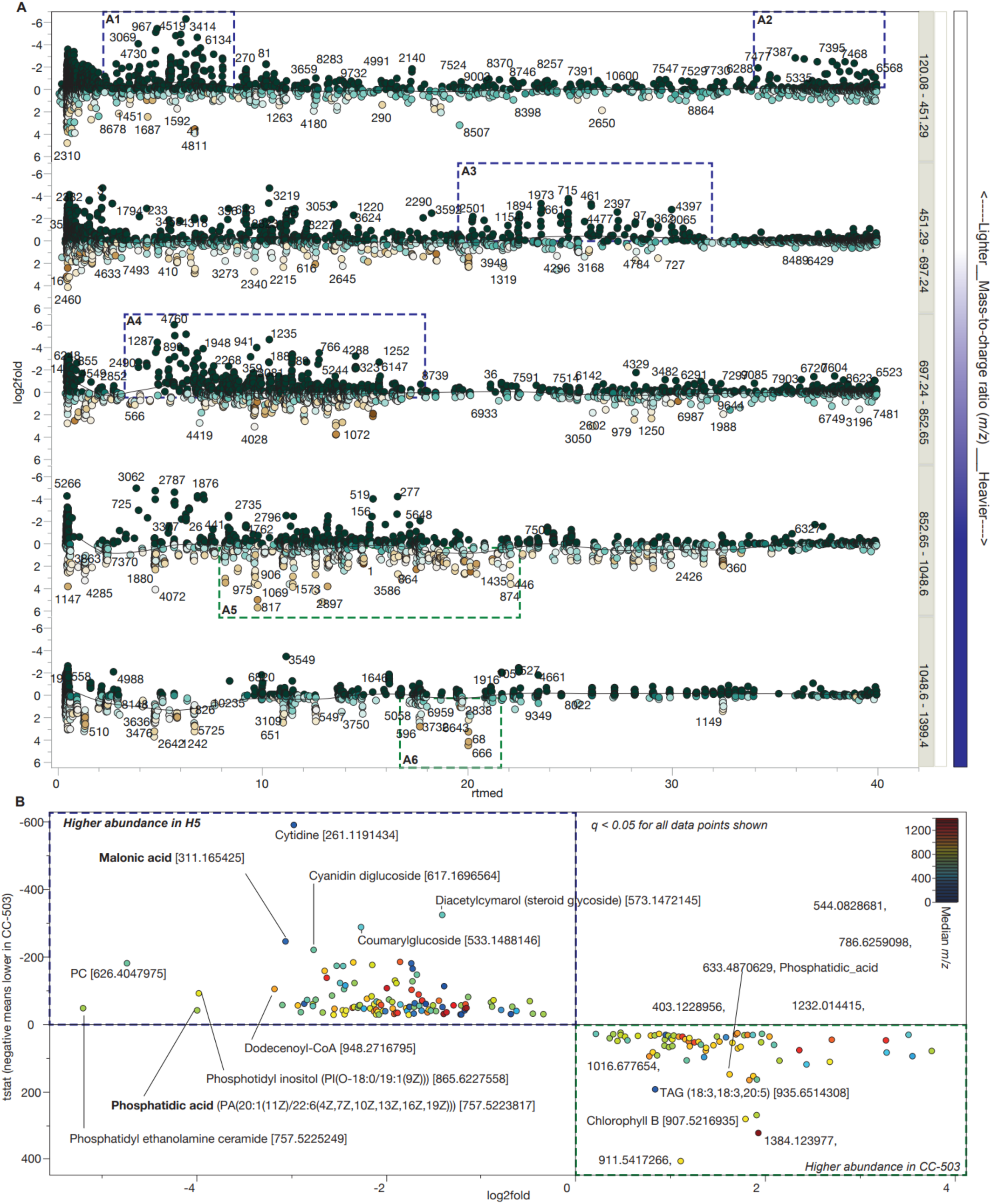
Metabolites detected in LC/MS-QToF metabolomic experiments. **(A)** Scatterplots are m/z bins showing comparative metabolomics for methanolic extracts of H5 vs. CC-503 at mid-log phase. This size bin shows an extensive array of various small, hydrophilic molecules accumulated in H5 (A1). Many free fatty acids were at higher concentrations in H5. In general, more compounds with later retention times in H5 indicate increased overall hydrophobicity of cell extracts. Accumulation of TAGs and free fatty acids in H5 explains the results described in (A2) and (A3; Glycerolipids, e.g., TAGs, increased in H5). Differential accumulation of H5 and CC-503 membrane lipids and their precursors in H5. High-impact mutations in enzymes in membrane lipid biosynthetic pathways are inferred to be responsible for the composition shift. Membrane lipids had lower diversity, while lipids containing saturated fatty acids were increased in H5 (A4-5). More high molecular weight (HMW) membrane lipids upregulated in CC-503 (A6). **(B)** Compound abundance (y-axis) and log 2-fold change (x-axis) in H5 compared to CC-503.

At Day 0, 21 transcripts expressed in CC-503 had no detectable expression in H5 (Table S3). Of these 21, three were annotated: HTR2, LCI1, and CGL16. The lack of high-temperature-responsive HTR2 suggests that H5 is less fit in the face of high-temperature stress. The low carbon-induced LCI1 protein [37] localizes to the plasma membrane and functions in active CO_2_ uptake. The CGL16 protein is conserved with unknown function, but knockouts of this gene in *C. reinhardtii* are metronidazole– and rose bengal-sensitive [38], indicating that its lack of expression in H5 may affect its hypoxic response. Of the three annotated genes without detectable expression in H5, LCI1 had the highest expression in CC-503 at an FPKM of 48.8. The lack of LCI1 expression is likely related to the upregulation of the carbonic anhydrases (CAH2/3) and low-carbon-induced (LCIC) protein in H5 (see Table S3).

In contrast, the periplasmic CAH2 was significantly upregulated (Table S3). The CAH1 and CAH2 genes are both α-type periplasmic enzymes, although the CAH1 protein is reportedly not expressed in CC-503 [42, 43]. Our results indicate that CAH1 the transcript is expressed in CC-503 (FPKM = 3,006.8) despite a repeat insertion preceding the TSS. These data reveal a scenario where mitochondrial CAH activity is lowered in H5 concurrent with upregulated periplasmic CAH activity. The β-CAHs 4/5 proteins are usually not made unless CO_2_ is limiting [44]. One interpretation of these results is that CO_2_ is less of a limiting factor for H5, possibly due to the upregulation of CAH2. Still, the high levels of malonate observed in our metabolomics analyses suggest that mitochondrial function is at least partially impaired in H5 and would be expected to suppress the expression of β-CAHs 4/5 proteins.

### Transcription factors (TFs) involved in lipid metabolism are significantly dysregulated in H5

We identified 23 putative transcription factors (TFs) during nutrient-replete growth in H5 from a total of 5,316 DEGs (2,832 upregulated genes and 2,484 down-regulated genes) using the Plant Transcription Factor Database (PlantTFDB) v4.0 [45] as a reference (Fig. 3, Table S3). We identified regulators for 43 upregulated and 39 down-regulated genes (Table S3). Fatty acid and TAG accumulation have been shown to increase following the overexpression of Dof-type [46] and MYB [47] TF families in *C. reinhardtii*. Basic helix-loop-helix (bHLH) TFs are stress response, cell growth, and division regulators in plants; overexpression of bHLH TFs enhances biomass and lipid production in the microalga *Nannochloropsis salina* [48]. Another regulator of lipid metabolism and stress tolerance that can regulate multiple genes under stress and enhance biomass and lipid productivity in microalgae (e.g., *Nannochloropsis salina*, [49]) is the bZip transcription factor. Four genes from the bHLH family were upregulated in H5 (2.8 – 4.1 – fold increase), and six genes from the bZip family were upregulated (2.8 – 22-fold increase). These results outline a drastic shift in the transcription factor landscape of H5 predominantly centered around cell growth and lipid accumulation pathways. The upregulation of a majority of transcription factors from bZip and bHLH families in H5 may positively affect cell growth due to their roles in light signaling and cell development while maintaining increased lipid accumulation [48, 49].

### H5 shows upregulated glycolysis leading to lipid accumulation

Many glycolytic enzymes exhibited significantly upregulated transcript expression in H5. These correspond with the high-impact mutation disrupting the regulatory region of 6-phosphofructokinase (PFK1, Enzyme Commission (EC) 2.7.1.11, Table S1) and may be fundamentally responsible for increased glycolysis in H5. The PFK1 enzyme is typically allosterically inhibited by ATP in an energy-balancing central regulator of glycolysis [50]. PFK1 activation correlates with the anoxic and mitochondrial inhibition responses. The activation of PFK1 in response to mitochondrial inhibition or hypoxia promotes glycolysis and starch degradation, also known as the Pasteur effect [51–53].

The upregulation of glycolysis likely fuels increased lipid biosynthesis in H5. The significant upregulation of pyruvate decarboxylase (PDC3, 1.18579-fold increase, *q =* 0.000210344, Table S3) supports this hypothesis. The PDC3 enzyme decarboxylates pyruvate from glycolysis for entry into fatty acid synthesis, which would ultimately have the effect of increased availability of substrate for lipid accumulation. The other key data supporting carbon-increased funneling from glycolysis into fatty acid production was the high levels of malonate observed in our metabolomic experiments. Other genes involved in starch and sucrose metabolism (Cre12.g488000, Cre06.g307150, Cre17.g721500, Cre12.g488050; KEGG pathway activation confidence score = 0.99712) and fructose and mannose metabolism (Cre10.g432900; KEGG score = 0.99755), as well as the pentose phosphate pathway (PPP; Cre03.g187450; KEGG score = 0.99265) were significantly upregulated in H5 in our KEGG analyses (Table S3). These metabolic enzymes normally have lower expression in nitrogen-limited conditions; their upregulation in H5 suggests intracellular nitrogen availability is not a primary factor in its enhanced lipid accumulation.

Five mitogen-activated protein kinases (MAPKs) were among the top 100 genes significantly upregulated in H5 (*q* < 0.05, Table S3). The function of this gene has yet to be experimentally verified, but other plant MAP kinases play roles in stress response and growth signaling. The numerous MAPKs upregulated in H5 suggest an influential role of these regulatory enzymes in the H5 phenotype. A 48 amino acid Parvulin-type peptidyl-prolyl cis-trans isomerase (PIN3, Cre07.g350400) was significantly upregulated in H5 in nitrogen-replete conditions (log 2-fold change = 2.18851*, q* = 0.00123182, Table S3). The PIN3 gene is a homolog of *Arabidopsis* auxin transporters.[39] Although the *C. reinhardtii* PIN3 is still uncharacterized, the mechanisms of auxin production in *C. reinhardtii* were elucidated in a recent preprint.[54] The authors found that auxin production from L-tryptophan is mediated by an extracellular enzyme L-amino acid oxidase (LAO1) [54]. If the function of this parvullin-like gene is conserved, H5 may have increased auxin uptake leading to accelerated growth even when undergoing a pseudo-nutrient deprivation response. Our evidence suggests, however, that this would be a downstream effect from the accelerated glycolysis resulting from the deregulated 6-phosphofructokinase (see also Table S1 and Fig. 3).

### H5 exhibits a cancer cell-like hypoxic nutrient deprivation response

Taken together with H5’s increased lipid production, our results outline a scenario wherein H5 displays a “hungry” tumor cell-like phenotype [55–57]. The disruption of several environmentally-responsive genes (e.g., betaine lipids) [58–61] also suggests a decrease of H5 in natural conditions, mirroring the honing towards higher productivity through loss of non-essential reactions commonly seen in microbial engineering (i.e., synthetic microbial chassis) [62].

Although H5 cells display expression patterns indicating accelerated glycolysis, they also have expression patterns of hypoxic cells (Table S3). Hypoxia-responsive genes upregulated in H5 included the hydrogenase maturation factor (HYDEF, 9.3-fold increase, *q =* 0.0002, Table S3), ferredoxin (FDX5, 21.4-fold increase, *q* = 0.0002), hybrid cluster protein 4 (HCP4, 7.4-fold increase, *q* = 0.0002, Table S3), and malate synthase (MAS1). The HCP4 hub gene was one of the most highly upregulated genes. It is highly upregulated under anoxic conditions and orchestrates *C. reinhardtii’s* anoxic response [63]. The FDX5 is linked to sulfur deprivation and hypoxic response networks in *C. reinhardtii* [64]. Without sulfur, *C. reinhardtii* cells cannot produce photogenic oxygen because they cannot support the rapid turnover of the photosystem II (PSII) D1 protein [65]. They eventually become hypoxic and can produce hydrogen (H_2_), among other biotechnologically interesting compounds [51, 66]. The expression of FDX5 is controlled by sulfur deprivation independently of anoxia, and its expression is negatively regulated by nitric oxide (NO) [65]. The upregulation of multiple hypoxia-responsive transcripts (Table S3) indicates that H5 experiences a pseudo-hypoxia state.

Four guanylate cyclases (CYGx) were significantly upregulated in H5 during exponential growth (Table S3). We predicted a loss-of-function frameshift mutation in the putative NO-producing guanylate cyclase Cre02.g100500 H5. A soluble guanylate cyclase (CYG56) mediates negative signaling by ammonium on nitrate reductase expression in *C. reinhardtii* [67]. The upregulation of the other four CYGs may be a compensatory mechanism for the disruption of Cre02.g100500. With the loss of NO production, FDX5 could be expected to increase in aerobic, sulfur-replete conditions, which we observed (Table S3). We investigated the possibility that the CYG Cre02.g100500 functions as an oxygen-sensing enzyme in *C. reinhardtii*; in this hypothesis, its ablation could explain the host of upregulated hypoxia-responsive genes in H5. However, the expression of Cre02.g100500 is extremely low in nutrient-replete conditions, precluding its possible role as a steady-state oxygen sensor. It is expressed only in nitrogen deprivation conditions in CC-503 (see https://conekt.sbs.ntu.edu.sg/), so its role in oxygen/nitrogen sensing is likely quite complex and dependent on multiple cofactors.

The copper deficiency responsive (CRD1) gene was significantly upregulated in H5 in nitrogen-replete conditions (6.7-fold increase, *q =* 0.0002, Table S3). This gene is usually expressed at trace levels in CC-503 and is only upregulated in response to extreme copper or oxygen deprivation [68]. The RSE1 gene was also significantly upregulated in H5 at Day 0 (10.5-fold increase, *q* = 0.00005, Table S3). Although without annotated function, RSE1 is highly responsive to copper starvation [69]. This copper-responsive expression of RSE1 led the authors to conclude that RSE1 is a likely target for the copper-response regulator CRR1. The copper starvation and hypoxia responses partially overlap in *C. reinhardtii* [68]; this overlapping set of genes was seen among the broad array of anoxia-responsive genes upregulated in H5. Malate synthase (MAS1) was significantly upregulated in H5 in nitrogen-replete conditions (4.2-fold increase, *q* = 0.0002, Table S3), corresponding to the up-regulation of MAS1 in response to nitrogen starvation and TAG accumulation seen in previous studies [70].

The ammonium transporter RHP1 was significantly upregulated in H5 at Day 0 (4.4-fold increase, *q* = 0.0002, Table S3). Combined, these results outline a nutrient deprivation response in H5. Tumor cells experience constant nutrient deprivation, and cancer cells have what is commonly referred to as a “hungry” phenotype [55–57]. Our multi-omics datasets for H5 suggest that it also experiences a “hungry” phenotype, facilitated by dysregulation of nutrient acquisition pathways, that drives its increased anabolism.

### The H5 nitrogen starvation response is unperturbed

In contrast to the nitrogen-replete H5 transcriptome, the nitrogen-deprived H5 transcriptome had few (number) genes expressed at significantly different levels than CC-503. Among these were the 443 amino acid low-CO2 inducible protein (LCIC). Its expression during nitrogen deprivation in H5 suggests that H5 experiences carbon deficiency when nitrogen deprived. A molybdate transporter (MOT2) was among the few genes with significantly different expression in H5 after three days of nitrogen starvation (Table S3). The *C. reinhardtii* molybdate transporter MOT1 is activated by nitrate, but MOT2 is only activated when extracellular molybdate concentrations are low [71]. This distinction supports the hypothesis that H5 is experiencing the deficiency of molybdenum, among other nutrients, in an exacerbated fashion compared to its wild-type progenitor. The iron-assimilatory protein FEA1 was significantly upregulated in H5 in nitrogen-replete and deprived conditions. The iron starvation response of *C. reinhardtii* is also known to induce LD and TAG formation [72]. In iron starvation, TAGs accumulate mainly from broken-down chloroplast membrane lipids [72]. One explanation for the drastic shift in lipid composition we observed is that chloroplast, but not endoplasmic reticulum (ER), lipids are broken down at a higher rate in H5. Metabolomics evidence corroborated this hypothesis, showing completely altered lipidomic profiles and increased phosphatidic acid in H5 (Data S1).

### Expression of photosynthetic machinery is upregulated in H5

We observed transcriptomic evidence that photosynthesis is upregulated in H5. In *C. reinhardtii*, the non-photochemical quenching (NPQ) of chlorophyll fluorescence (qE) depends on light-induced accumulation of the LHCSR proteins, specifically LHCSR3 [73] In contrast, under the absence of LHCSR proteins, the structure of the PSII-LHCII supercomplexes changes in the HL-acclimated state to shift the distribution of the PSII antenna protein LHCBM5 toward the PSII core-enriched fraction B3 [74]. The primary LHCII complex forms from nine genes in *C. reinhardtii,* which group based on their sequence homology: Type I (LHCBM3, LHCBM4, LHCBM6, LHCBM8, LHCBM9), Type II (LHCBM5), Type III (LHCBM2, LHCBM7) and Type IV (LHCBM1).[75] In this study, five out of nine LHCII complex genes were upregulated in H5. This change is consistent with the alteration of the qE trigger mechanism from the LHCSR3 to the LHCII complex. The lack of LHCBM1 can reduce qE [76] and is needed to develop high NPQ as a partner for LHCSR3 [77].

A sulfite oxidase (SUOX1) was significantly upregulated in H5 in nitrogen-replete conditions. This enzyme recycles intracellular sulfur in *C. reinhardtii*. The increased accumulation of the SUOX transcript during sulfur deprivation implies that H5 breaks down intracellular sulfur molecules more rapidly than CC-503, again mirroring a nutrient deprivation response or ‘hungry’ state commonly seen in cancer cells [55–57]. The necessity may be from the higher turnover of the D1 protein of photosystem II (PSII). Eight of nine PSII (PsbX) proteins with significantly different expression in H5 were up-regulated (Table S3), indicating that photosynthesis is generally upregulated in H5.

### Glycolytic and lipidomic shifts underlie the re-routing of H5 intracellular carbon to TAGs

Metabolites from cell pellets of H5 and CC-503 from the exponential growth phase (OD_600_ = 0.4; i.e., nutrient-replete conditions) underwent microwave-assisted extraction with methanol [78, 79]. Metabolites were loaded on a C-18 reverse phase column (Agilent, Santa Clara, CA, USA) and eluted using a semi-linear isopropanol gradient. Eluted compounds were interrogated with a QToF mass spectrometer (Fig 4, Data S1). The resultant mass-to-charge ratio (*m/z*), ion abundance, and retention times were analyzed using the XCMS software suite (https://xcmsonline.scripps.edu) using nonlinear peak alignment, matching, and identification.

Overall, 7,692 molecular features were detected in CC-503 and H5 methanolic extracts. Of these, 2,196 had significant differences in abundance between the two strains (*p* < 0.01). Using XCMS metabolomics screening, we identified 5,168 compounds correlating with the detected molecular features with differential abundance (*p* < 0.00001). Notably, we observed several metabolites with roles in lipid accumulation with differential abundance in CC-503 and the H5 mutant (Fig. 4). Compounds with putative signaling roles, including cyclopentadione and dihydrostilbene derivatives, were present at elevated levels or uniquely present in H5 (Fig. 4, Data S1). Pentadiones and stilbenes are hormone-like compounds that modulate energy metabolism [80, 81]. We note that the unique presence of these molecules correlated with a high-impact mutation in a prostaglandin-F synthase (EC1.1.1.188, Data S1). Additionally, ethanamine, a nitric oxide donor [82], was more abundant in H5 (Data S1). The dysregulation of these molecules hint at disrupted hormonal signaling in H5.

We found 12 intermediate metabolites in the glycolipid and 11 in the glycerolipid desaturation pathways significantly dysregulated (*p* < 0.05) in H5. Malonate, an inhibitor of oxidative phosphorylation and promotor of glycolysis and cell proliferation, was ∼10x higher in H5 (*p* = 8.5 x 10 ^-4^) while chlorophyll *a* and *b* concentrations were 2.3– and 3.5-fold higher in CC-503 (Data S1 i.e., HPLC-MS metabolomics datasets). A gene coding for a glycerol kinase had a frameshift mutation in H5 (Table S1). As glycerol kinase catalyzes the phosphorylation of glycerol to form glycerol 3-phosphate, a component of glycerophospholipids, the loss-of-function mutation in a glycerol kinase would change the composition of lipids by decreasing the phospholipid population. A mutation in a lipase was also seen (Table S1). A loss-of-function mutation in this lipase would prevent the formation of product DAGs. Our metabolomics data shows a radical shift in all high-molecular-weight lipid populations, including TAGs, DAGs, and monoacylglycerols (MAGs). Many diacylglycerols (DAGs) were at lower concentrations in H5; however, the diversity of triacylglycerol classes increased. Nine TAG moieties were increased in H5, while five were decreased. Diacylglycerols (DAGs) were reduced in H5 compared to CC-503. The metabolomic and transcriptomic evidence suggests that TAG to DAG formation is perturbed in H5. Decreases in MGDG have coincided with increased TAG accumulation [72]. The higher diversity of TAGs may directly result from more energy and carbon allocation into TAG production after eliminating non-essential, conditional, standby metabolisms.

Betaine lipids are crucial for the adaptation to phosphate stress in some microalgae [83]. We observed four out of five detectable betaine lipids downregulated in H5 (Data S1). The lipids DGTA(18:1/22:4(10Z,13Z,16Z,19Z), DGTA(18:1/22:4(10Z,13Z,16Z,19Z)), DGTA(18:1/22:4(10Z,13Z,16Z,19Z)), and DGTS(16:0/16:0) were higher in CC-503, while DGTS(16:0/18:2(9Z,12Z)) was higher in H5 (Data S1). In addition to indicating a possible phosphate deficiency surplus, the absence of betaine lipids might be directly related to overall lipid accumulation in H5. Betaine is an efficient methyl donor known to increase nitrogen storage capacity and promote fat mobilization, as well as changing cellular fat content and composition [84]. Thus, the lack of betaine lipids in H5 likely has an influential role in its metabolism.

We observed very high levels of the nucleotide cytidine in H5 compared to CC-503 (Data S1). Abnormally high intracellular nucleotide levels and their metabolic demand are characteristic of tumor cells [85]; here, the increased cytidine appears to result from a block in its condensation with DAGs. The expression of a cytidine diphosphate diacylglycerol synthase (CDP-DAG) was down-regulated in H5 (Table S3, Fig. S3), corresponding with the absence of CDP-DAGs and high levels of intracellular cytidine (Data S1). These results suggest that the condensation of cytidine with phosphatic acid [86] is blocked in H5, which would be expected to reroute carbon flux into TAG production.

Malonate was also increased in H5, which indicates a strongly positive cellular energy balance. Malonate is an essential precursor and rate-limiting reactant in fatty acid biosynthesis; thus, its increased availability suggests a potential for higher lipid production. Excess malonate is toxic to mitochondria [87]. It inhibits succinate dehydrogenase, disrupting the electron transport chain driving oxidative phosphorylation, and diverting metabolism into glycolysis [88] Increased cell proliferation is also an effect of increased malonate; the high accumulation of this compound in H5 explains how H5 accumulates lipids while still proliferating at wild-type-like growth rates.

Malonate accumulation is also observed in acute myeloid leukemia cells, corresponding to higher pyruvate carboxylase activity [89]. We note that The H5 pyruvate carboxylase PC3 was significantly upregulated compared to CC-503, indicating its likely role in facilitating this route of CO_2_ sequestration into lipids. The significantly upregulated H5 carbonic anhydrase CAH2 hints at a possible entry valve opening to allow for greater carbon intake for a hungrier *C. reinhardtii* cell that is deficient in key lipid breakdown and accessory metabolic pathways.

### Whole genome bisulfite sequencing reveals key epigenomic differences in H5

To test if H5 cells have different methylation patterns than the parental cells, we carried out whole genome bisulfite sequencing on CC-503 and H5 in nutrient-replete conditions (Data S2). Correcting for multiple hypotheses, ANOVA analysis of the of 250 bp scanning windows of H5 genome showed significant hyper methylation in 14,720 transcribed regions compared to CC-503. Of these, 9,599 were hypomethylated, and 5,121 were hypermethylated. Among significantly hypomethylated genes were Activin receptor-Like Kinase (ALK, [90, 91] methylation difference (MtD) = –0.026587384, *q* = 0.015702734), transforming growth factor beta (TGF-beta) type I receptor predominantly expressed in actively proliferating cells [92–95], and the high-level microRNA regulator RabGAP/TBC CGL44 (MtD = –0.021276596, *q* = 0.00363009) whose function was predicted by ChlamyNET [96] as GTPase activity (GO:0043087)) associated with TAG production.

Among significantly hypermethylated genes was the mitochondrial alternative oxidase (AOX1, MtD = 0.018961256, *q* = 0.042157736, Data S2). Changes in this gene concur with our transcriptomics (Table S3) and metabolomics (Data S1) analyses that also suggest mitochondrial dysfunction in H5. Notably, the cofactor of nitrate reductase and xanthine dehydrogenaase 3 (CNX3) was hypermethylated in H5 (MtD = 0.042133865, *q* = 0.010310439, Data S2). This gene encodes a protein involved in molybdenum cofactor biosynthesis. We note that this observed significant hypermethylation correlates with the increased expression of the molybdenum transporter MOT2 in H5 in nutrient-replete and nitrogen-deprived conditions (Fig. 3 and Table S3). Their co-occurrence indicates that molybdenum metabolism is substantially perturbed in H5.

## CONCLUSIONS

Our multi-omic analyses reveal that the high-lipid phenotype of strain H5 is driven by a metabolic shift reminiscent of cancer cell metabolism. H5 exhibits hallmark features such as upregulated glycolysis and pseudo-hypoxia, which correlate with a dramatic accumulation of malonate—a key metabolite known to arise from glycolytic flux and to inhibit mitochondrial function while promoting lipid storage (Fig. 5).

**Figure 5.**
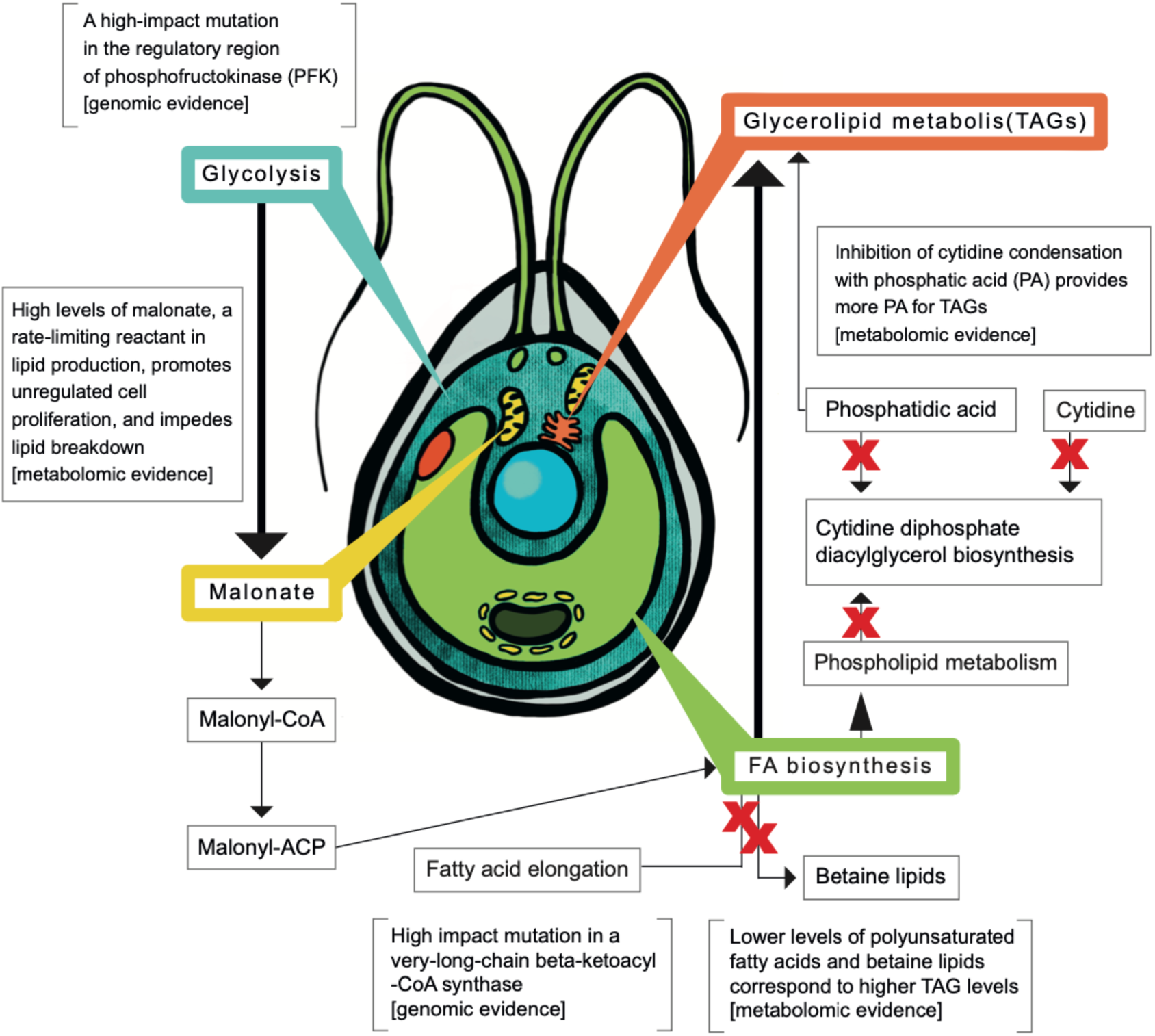
Overview of multi-omics evidence explaining the H5 phenotype. The mutant strain H5 accumulates lipids during exponential phase growth through a combination of enhanced glycolysis and rerouting carbon from non-essential pathways to TAG and membrane lipid biosynthesis.

A disruptive mutation in the regulatory domain of 6-phosphofructokinase (PFK; EC 2.7.1.11) in H5 may weaken ATP-mediated feedback inhibition, effectively derepressing glycolysis. This mutation is supported by increased malonate levels (Table S3, Data S1), and the CLiP PFK mutant (PfkA, Cre11.g467552) is among the strongest lipid accumulators, pointing to this alteration as a central driver of the phenotype. In addition to the PFK mutation, H5 carries other synergistic mutations that mimic nutrient stress and streamline carbon flow into lipid biosynthesis. This deregulated, cancer-like metabolism—though pathological in multicellular organisms—is beneficial in controlled lab conditions where biomass and lipid output are prioritized.

The loss of non-essential pathways following UV mutagenesis appears to enhance resource allocation toward lipid production. Just as a sculptor reveals form by removing the superfluous, artificial evolution can pare down cellular complexity to yield optimized microbial chassis for industrial use.

Finally, the parallels between tumor metabolism and lipid-producing algae underscore glycolysis, acetyl-CoA flux, and de novo lipogenesis as valuable engineering targets. While targeted metabolic engineering offers a rational path forward, artificial selection may ultimately uncover more robust, emergent solutions. These findings open the door to novel bioengineering strategies for creating high-yield strains for sustainable biofuel and bioproduct generation.

## METHODS

### Algal strains, culture conditions, and growth analysis

*C. reinhardtii* strain CC-503 was purchased from the *Chlamydomonas* Resource Center (http://Chlamycollection.org), and H5 was generated from our previous study [17]. We deposited H5 in the *Chlamydomonas* Resource Center (CC-5163 H5-M1 mt+ (high lipid-producing strain)) (https://www.Chlamycollection.org/product/cc-5163-h5-m1-mt-high-lipid-producing/). *C. reinhardtii* cultures were grown on tris-acetate-phosphate [15] medium using constant lights with an intensity of 400 μmol photons m^−2^ s^−1^. The pond simulators PBR101 (Phenometrics, Inc., Michigan, USA) were approximately cylinder photobioreactors (PBRs). They were set up with a working volume of 360 mL and a light depth of 14.0 cm for each working PBR. The PBRs were injected with air enriched with 1.0% CO_2_ at a flow rate of 0.36 L/min. Temperature and pH were maintained at 25 ± 1 °C and 7.0 ± 0.5, respectively. Stirring was set at a constant speed of 100 RPM using a 28.6 mm stir bar. Cell diameter and size distribution were determined using the Cellometer Auto M10 (Nexcelom Bioscience LLC, Lawrence, MA, USA). The growth and FACS were carried out on H5 and CC-503 strains obtained from the *Chlamydomonas* Resource Center (https://www.Chlamycollection.org, St. Paul, MN, USA).

### Analysis of lipid accumulation in different mutant strains using a fluorescent spectrophotometer

A group of 35 strains was purchased from *Chlamydomonas* Resource Center (http://Chlamycollection.org). These strains were sub-cultured several times to reach exponential growth in 15 ml Falcon™ tubes. The freshly prepared cultures from the exponential growth phase were used as seed cultures and transferred to 96-well microplates with a volume of 200 μL for each well and a starting OD_600_ of 0.1. After these cultures reached the exponential growth phase (OD_600_ ∼ 0.4) in 96-well microplates, cells were stained with 8.0 μg/ml BODIPY 505/515 for 15 min. The cell pellets were then collected by centrifugation, washed, and resuspended with the same volume of TAP medium. The fluorescence of stained cells was determined with an excitation wavelength of 485 nm and an emission wavelength of 515 nm using the hybrid multimode microplate reader Synergy H1 (BioTek Instruments, Winooski, Vermont, United States), according to the manufacturer.

### UV mutagenesis

The H5 mutant was generated in a previous study [17]. To summarize, *C. reinhardtii* (strain CC-503) cells were exposed to ultraviolet (UV) light at a distance of 30 cm (253.7 nm,100 μW/cm^2^, 60 HZ, NuAire, Plymouth, MN, USA) for 2 min, then kept in the dark for 24 hours to suppress DNA repair. Empirical pilot studies helped to identify a UV dosage that gave ∼5% survival in a lawn of approximately 1 billion cells on a 100×15mm petri dish. Every round of mutagenesis was performed at this optimal dosage.

The sorted cells were grown under light in a liquid tris-acetate-phosphate (TAP) medium [97], and the same process was applied to the third and fourth rounds [16, 17]. The cells selected after the fourth round were sorted again without mutagenesis to enrich the lipid-accumulating cells further. The lipophilic dye BODIPY505/515 was used to label high-lipid mutants due to its high oil/water partition coefficient, enabling it to permeate membranes to distinctively label neutral lipids *in vivo* [98]. Stained cell population sorting by FACS partitioned sub-populations enriched for lipid production.

Aliquots of cells from the resulting colonies were stained by BODIPY to illuminate cells containing neutral lipids that were then sorted using a FACSaria BDIII cell sorter (BD Biosciences-Becton, Dickinson, and Company, San Jose, CA). Cells passing multiple gates, including high BODIPY fluorescence, were collected and grown for three more rounds of selection, totaling four rounds of UV mutagenesis and FACS sorting [16]. Two additional rounds of selection were carried out to further enrich for lipid accumulating cells. CC-503 and the single colony (the mutant H5 strain) that produced the highest signal were selected for all future analyses, including genome and transcriptome sequencing. Analyses were performed on the strain directly after its ‘high-lipid’ characterization (i.e., at 3.2-fold more lipids than its parental CC-503 quantified by BODIPY fluorescence). We note that after two years of maintenance in our laboratory, the H5 mutant lipid accumulation declined from 3.2 to 1.5-fold relative to CC-503. However, its wild-type-like growth rate was maintained (Fig. S1). The reversion may be due to subsequent epigenetic changes in the H5 strain or the accumulation of spontaneous mutations that suppress lipid accumulation.

### Generation of genome sequence and assembly of H5 and variant detection

CC-503 and the H5 mutant single colony that produced the highest FACS signal were selected for genome sequencing. Genomic PCR-free sequencing of H5 by HISeq2500 and SNP calling was run using Phytozome version 10.3 as the reference. We used Bowtie2 [99] to align the reads with the reference genome PhytozomeV10/Creinhardtii/Creinhardtii_281_v5.0.fa. Candidate SNPs were identified using SAMtools [100]. SnpEff [24] was used to annotate detected variants with default parameters using *C. reinhardtii*_281_v5.0 as a database. We focused on ‘high-impact’ mutations for phenotype attributions. The scale of impact in SnpEff was measured as the ‘effect’ variable (EFF) where EFF= Effect (Effect_Impact | Functional_Class | Codon_Change | Amino_Acid_Change| Amino_Acid_Length | Gene_Name | Transcript_BioType | Gene_Coding | Transcript_ID | Exon_Rank | Genotype_Number) and the multiple effects comprise the weight multiplier formula for SnpEff variant calling.

### Transcriptomic analysis of wild-type CC-503 and H5 mutant

The genome sequence of *C. reinhardtii* was used as a reference to align the transcriptome reads. Using Tuxedo suite, which includes Bowtie2 [99, 101], TopHat (https://github.com/DaehwanKimLab/tophat), and Cufflinks (http://cole-trapnell-lab.github.io/cufflinks/), the raw reads were aligned to the *C. reinhardtii* reference genome (downloaded from http://genome.jgi.doe.gov/). For differentially expressed genes, FPKM values (Fragments Per Kilobase of transcript per Million mapped reads) [102] were calculated, a normalized measure of reading density that allows transcript levels to be compared within and between samples quantitatively.

Transcriptomic sequencing (RNAseq) for CC-503 and H5 cultures in triplicates was performed at 0, 1, and 3 days of nitrogen starvation (performed as in Wang et al., [34]; see Table S3). Mid-log phase (OD_600_ = 0.4) cells were harvested for the Day 0 timepoint. To define the contributions of deprived and nutrient-replete states towards lipid accumulation in H5 and CC-503, transcriptome analyses were performed at zero, one, and three days post-inoculation in Cuffdiff[35] (Tables S2 and S3). The differentially expressed genes (DEGs) were screened at false discovery rate (FDR) –corrected *p* values (i.e., *q* values) ≤ 0.05 (Table S3). At Day 0, 11,126 and 10,554 genes were expressed in CC-503 and H5 at FPKM > 1.0. Of these expressed genes, 10,094 were shared between the two strains, 1,032 were unique in CC-503, and 460 were unique in H5. Differentially expressed genes from Day 0 (starting point, non-starvation control) and Day 1 (deprived for 24 H) comprised 1,798 and 1,727 upregulated transcripts and 1,296 and 1,602 down-regulated transcripts.

The algorithm Cuffdiff [35] was used to identify differentially expressed genes (DEGs) between two groups of samples (triplicates for each group, see Table S3). Functional and gene set enrichment analysis of DEGs was performed using BiNGO [103] plugin version 3.0.3 in Cytoscape. BiNGO determined the statistical overrepresentation of the DEGs genes over the Gene Ontology (GO) terms. The *p* values were calculated using the hypergeometric test, and Benjamini and Hochberg False Discovery Rate (FDR) correction [104] was used to identify the statistical significance of gene ontology terms with corrected *p(q)* < 0.05 for multiple testing [103].

Transcript-level functional annotations were assigned using a custom pipeline developed in our prior work on algal metabolic network model refinement [102]. Briefly, predicted proteins were functionally annotated using BLAST2GO as in [105], integrating BLASTP homology searches, InterProScan domain identification, and Gene Ontology (GO) term assignment. This approach provided a broad functional context based on sequence similarity to well-characterized protein families and domains.

For pathway enrichment analysis, we used the Algal Functional Annotation Tool (AFAT, http://pathways.mcdb.ucla.edu/algal/) [106], which is specifically designed for analyzing photosynthetic eukaryotic genomes. Importantly, AFAT relies exclusively on Phytozome v5 gene accessions as its reference background, including *C. reinhardtii* and other green algae. Thus, all enrichment analyses were constrained to plant– and algae-specific orthologs, ensuring taxonomic consistency.

GO term and KEGG pathway enrichments were computed using Fisher’s Exact Test with Benjamini-Hochberg correction (FDR < 0.05). KEGG pathway identifiers are based on standard KEGG nomenclature, which can result in pathway names that reference non-photosynthetic organisms (e.g., Vibrio cholerae infection). These labels reflect the historical origins of pathway discovery, not the taxonomic origin of genes in our dataset. For example, conserved eukaryotic processes such as vesicle trafficking and protein transport may appear under SNARE or antigen-processing pathways due to shared components, even though the genes are *Chlamydomonas*-derived.

This annotation strategy mirrors established practice in algal transcriptomic studies [105] where mixed-lineage pathway labels appear because of cross-kingdom conservation of molecular functions. To prevent confusion, we explicitly note that all gene IDs used in enrichment analyses were *C. reinhardtii* –specific, and no foreign or bacterial genes were present in our assemblies or expression sets.

### Bisulfite sequencing and methylation analyses

DNA was extracted from three H5 and two CC-503 colonies (as control) using the DNeasy Plant Mini Kit. (Qiagen, Hilden, Germany). Libraries were prepared using the Zymo EZ DNA Methylation Gold Kit (Zymo Research, CA, USA) and Swift Accel-NGS MethylSeq (Swift Biosciences, Inc, MI, USA). Whole Genome Bisulfite Sequencing and library preparation were carried out at Admera Health (South Plainfield, NJ, USA). For each sample, approximately 31 million 2 x 150 paired end reads were obtained. Methylation analysis was done using the bisulfate analysis package of CLC Genomics Workbench V.22 (QIAGEN, Aarhus, Denmark). Fisher’s Exact Test was used to examine the statistical significance of differential methylation in CHG context (H=A, C, or T) of 100-base frames between the control and H5 reads using annotated *C. reinhardtii* transcripts (https://phytozome-next.jgi.doe.gov/info/Creinhardtii_v5_6).

### Confocal microscope visualization of CC-503 and H5 mutant

CC-503 and H5 mutant from FACS selection after the fourth round of UV mutagenesis were grown for three days at 25 °C in a Photon System Instrument (PSI) growth chamber (AlgaeTron, Drasov, Czech Republic) in both standard TAP liquid and nitrogen-deficient media. Cell optical density was measured at 600 nm using a microplate reader (Synergy H1, microplate reader, BioTek, Singapore) and normalized for staining. Cells were observed with an Olympus (Tokyo, Japan) Fluoview 1000 confocal laser-scanning microscope using 405, 488, 559, and 635 nm lasers to visualize BODIPY 505/515 (Schenectady, NY, USA) stained cells. Objective: 60x NA 1.3, Laser power, 1 uW for GFP (Filter: 515 – 560 nm) and chlorophyll autofluorescence (590-650nm), The Pixel size is 0.041um (10x).

For BODIPY staining, the 488 nm excitation laser was used. A photomultiplier tube (PMT) detector was used to collect the light, and the spectral filter was set from 520 to 580 nm. Lipid staining was performed as follows:1 μL of 10 mg/mL BODIPY 505/515 was added to 100 μL of cells (approximately 10^7^ cells/mL) and allowed to incubate at room temperature for one hour. If fluorescence quenching was too rapid to allow for imaging, cells were incubated for more extended periods before microscopy.

For the extraction, cultured microalgae were scraped from agar plates after one week growth under approximately 100 μmol·m^-2^ ·s^-1^ and equal weights of three replicates for each strain were placed into five mL methanol then vortexed. Algal-methanol solutions were microwaved on high power (1150 watts, 2450 Mhz) in a ME732K microwave (Samsung, Suwon-si, South Korea) with a triple distribution system. Solutions were microwaved until boiling five times and then vortexed. Extracts were filtered with a 5 µm then 2-µm filter (Millipore, Billerica, MA, USA) and maintained in the dark at 4 °C.

The HPLC method was developed in-house and is based on a comprehensive shotgun lipidome technique [107]. Extracted metabolites were loaded on a reverse-phase C18 column starting with an elution buffer composition of 36% 30 mM ammonium formate (AF, HCO2NH4, Chemical Abstract Services number (CAS): 540-69-2, MW: 63.06. EC Number: 208-753-9; Sigma-Aldrich, Darmstadt, DE), 36% acetonitrile (CAN, CH3CN, Purity 99.8%, Molecular Weight = 41.05, CAS: 50-165-7178; Sigma-Aldrich, Darmstadt, DE), and 28% isopropanol (IPA, (CH3)2CHOH, MW: 60.10, purity ≥99.5%, CAS: 67-63-0; Sigma-Aldrich, Darmstadt, DE). A semi-linear gradient was applied to the column resulting in 90% isopropanol and 10% acetonitrile for 18 min at 300 μL/min and with a column temperature of 50°C on an Agilent 1290 Infinity II LCMS with a UHPLC column and the stream was diverted to a quadrupole time-of-flight mass spectrometer (Agilent LCMS QToF 6538, Agilent, Santa Clara, USA). Accurate mass profiling was enabled by lock mass compounds at 121.050873 and 922.009798 m/z. Collision energies were assigned based on the formula [1.4*(m/z)/100) + 20] developed to maintain small molecule integrity and fragment larger ions.

Source datasets are in Data S1 and are comprised of high-performance liquid chromatography (HPLC) and quadrupole time-of-flight mass spectrometry (MS-QTOF, positive polarity mode) XCMS[108–111] analyses. Compound identification and comparisons were performed using nonlinear peak alignment and matching against existing databases (XCMS [108–111], HMDB [112]). This dataset includes ion abundances, total ion chromatograms (TICs)), –fold and log 2-fold change values, compound retention times, T-values, *p* values, and *q* values (*p* values adjusted for false discovery rates). Also included in the.zip file are extracted ion chromatograms (EICs) for individual compounds and boxplots of each compound compared between CC-503 and H5.

### Computational metabolomics analyses

Compounds identified with HPLC-MS/QToF were processed in the XCMS online software suite (https://xcmsonline.scripps.edu/. The XCMS suite integrates Lipid Maps (http://www.lipidmaps.org/), Kyoto Encylopedia for Genes and Genomes (KEGG; http://www.kegg.jp/kegg/kegg2.html), Human Metabolome Database (HMDB; http://www.hmdb.ca/), Scripps Center for Metabolomics (https://metlin.scripps.edu/index.php), the National Institute for Standards and Technology (NIST; http://webbook.nist.gov/chemistry/), and Massbank (http://www.massbank.jp) to provide a comprehensive service for identifying molecules from mass spectrometry data and comparing them between samples. The XCMS suite uses multiple pathway-based analysis tools to identify metabolic pathways from collections of candidate models, compare them between samples, and derive statistically significant results for altered levels of metabolites and pathway activity between sample groups.

Mummichog (https://shuzhao-li.github.io/mummichog.org/) was able to resolve differences in metabolic pathways for lipid metabolism. The Mummichog (1.1.6) command was: [mummichog/main.py –f H5.vs.CC.503.tsv –r ref-list.txt –k./temp –o mummichog –m positive –n A1000047 –u 5 –c 0 –g 0]. The *C. reinhardtii* metabolic network was loaded for feature searches (BioCyc6.0). Overall, 1,617 input compounds were used as queries to find 567 Mummichog compounds (518 at *p* < 0.005) from 11,892 references. Resampling was done with 100 permutations to estimate the background, and the pathway background was estimated on 33,200 random pathway values. The null distribution was estimated on 807 random modules, and our data matched seven network modules (see also Data S1).

### Graphics

Fig. 1 was made using Adobe Illustrator and Photoshop, Microsoft Excel, and the Olympus (Tokyo, Japan) CellSens software. Fig. 2 was made using Adobe Photoshop, SnpEff [24], Circosviz (https://github.com/MadsAlbertsen/multi-metagenome/blob/master/circosviz/circosviz.pl), and Circos (circos.ca). Figs. 3 and 4 were made in R (3.x) and Adobe Illustrator.

### Statistical analyses

Biological triplicates (i.e., H5 cell populations grown from three independent colonies) were analyzed for RNAseq mapping counts to determine up– and down-regulated genes in H5 (Table S3). A student’s *t* test was performed to evaluate the significance of the difference between the two groups of samples for growth data (*p* < 0.05). Our LC/MS-qTOF results were significant at *p* < 0.05 after correction for multiple hypotheses (*q* values). Statistical metrics are included in the LC/MS-qTOF supplementary dataset describing retention time corrections and false-discovery rate-corrected *p* values (*q* values) [108] for matches against databases other than CHLAMY BioCyc6.

## DATA AVAILABILITY STATEMENT

Supplementary data are available at Zenodo DOI: https://doi.org/10.5281/zenodo.14525303. The repository includes raw metabolomics data—total ion chromatograms (TICs) and extracted ion chromatograms (EICs)—as well as all supplementary datasets associated with the analyses. Raw sequencing data for the omics experiments are deposited in the NCBI SRA under the following Bioprojects: PRJNA1216387 (genomic reads for variant analysis), PRJNA1216608 and PRJNA389185 (RNA-seq for nitrogen starvation transcriptomics), and PRJNA1158323 (whole-genome bisulfite sequencing for epigenomics).

## DECLARATION OF GENERATIVE AT AND AI-ASSISTED TECHNOLOGIES IN THE WRITING PROCESS

During the preparation of this work the authors used OpenAI’s ChatGPT in order to condense otherwise verbose writing. After using this tool/service, the authors reviewed and edited the content as needed and take full responsibility for the content of the published article.

## ACKNOWLEDGEMENTS

Financial support for this work was provided by New York University Abu Dhabi Faculty Research Funds (AD060) and by the NYUAD Institute grant (G1205-1205i). We thank Marc Arnoux and the CGSB Sequencing Core for high throughput sequencing, Nizar Drou and CGSB Bioinformatics Core for raw RNAseq data processing, and NYUAD High-Performance Computing and Core Technology Platforms for their resources and logistical support.

## AUTHOR CONTRIBUTIONS

D.R.N., A.C., W.F., K.S.-A, designed the study. D.R.N., A.C., A.J., B.D., K.M.H, K.S.-A., and L.N. carried out computational analyses, D.R.N., A.C., W.F., K.S.-A., B.K., B.D., A.J., D.A.-K., A.M., L.A.N., L.N., S.D., A.S.A., carried out growth and morphometric studies, A.M., L.N., generated samples for bisulfate analysis, M. J. O’C. carried out LCMS. M.S., carried out confocal microscopy, all authors contributed to the critical reviewing and editing of the manuscript; K.S.-A. supervised the study.

